# Transgenic *Drosophila* lines for LexA-dependent gene and growth regulation

**DOI:** 10.1101/2021.10.07.463520

**Authors:** Kathleen R. Chang, Deborah D. Tsao, Celine Bennett, Elaine Wang, Jax F. Floyd, Ashley S.Y. Tay, Emily Greenwald, Ella S. Kim, Catherine Griffin, Elizabeth Morse, Townley Chisholm, Anne E. Rankin, Alberto Baena-Lopez, Nicole Lantz, Elizabeth Fox, Lutz Kockel, Seung K. Kim, Sangbin Park

**Affiliations:** Stanford University, Stanford, CA 94305; Dept. of Developmental Biology, Stanford University School of Medicine, Stanford CA 94305; The Lawrenceville School, 2500 Main St, NJ 08648; Department of Genetics, Stanford University School of Medicine, Stanford CA 94305; Phillips Exeter Academy, Exeter, NH 03833; Dunn School of Pathology, Oxford University, Oxford, U.K.; Departments of Medicine and of Pediatrics, Stanford University School of Medicine, Stanford, CA 94305; Stanford Diabetes Research Center, Stanford, CA 94305

**Author notes:** These authors contributed equally (Celine Bennett, Elaine Wang, Jax F. Floyd, Ashley S.Y. Tay).

## Abstract

Conditional expression of short hairpin RNAs (shRNAs) with binary genetic systems is an indispensable tool for studying gene function. Addressing mechanisms underlying cell-cell communication *in vivo* benefits from simultaneous use of two independent gene expression systems. To complement the abundance of existing Gal4/UAS-based resources in *Drosophila*, we and others have developed LexA/LexAop-based genetic tools. Here, we describe experimental and pedagogical advances that promote the efficient conversion of *Drosophila* Gal4 lines to LexA lines, and the generation of LexAop-shRNA lines to suppress gene function. We developed a CRISPR/Cas9-based knock-in system to replace Gal4 coding sequences with LexA, and a LexAop-based shRNA expression vector to achieve shRNA-mediated gene silencing. We demonstrate the use of these approaches to achieve targeted genetic loss-of-function in multiple tissues. We also detail our development of secondary school curricula that enable students to create transgenic flies, thereby magnifying the production of well-characterized LexA/LexAop lines for the scientific community. The genetic tools and teaching methods presented here provide LexA/LexAop resources that complement existing resources to study intercellular communication coordinating metazoan physiology and development.

## INTRODUCTION

Binary gene expression systems are a cornerstone in *Drosophila* investigations of gene regulation and function. The most widely-used binary expression system in *Drosophila* relies on two separate components: a yeast Gal4-based transcriptional transactivator whose expression is under control of cell-type specific enhancers, and Gal4-responsive upstream activating sequence (UAS) that regulates the expression of target genes (Brand and Perrimon 1993; Hayashi et al 2002; Gohl et al 2011). To investigate the impact of targeted gene suppression, investigators have used Gal4/UAS-mediated gene knockdown to express short hairpin RNAs (shRNAs) in specific cell types (Ni et al 2011). Studies of many biological questions require simultaneous manipulation of two or more independent cell populations or genes (reviewed in Kim et al 2021). These studies often benefit from multiple binary systems combined in a single fly to study genetic perturbations in multiple tissues. Such a multiplex approach requires systems that function independently of the UAS/Gal4 system, such as the LexA/LexAop binary system, comprised of a bacterial transactivator (LexA) expressed from cell-type specific *Drosophila* enhancers and a *sulA*-derived LexA operator-promoter (LexAop) that drives the expression of adjacent target genes (Pfeiffer et al 2010).

To address this need, we and others have made systematic efforts to expand the number of well-characterized, publicly available LexA fly lines, either by linking defined enhancers to sequences encoding LexA (Pfeiffer et al 2010) or by mobilizing a LexA-containing transposable element to insert at genomic sites near enhancers, resulting in ‘enhancer trap’ LexA lines with unique expression patterns (Kockel et al 2016, 2019). Although these resources are slowly growing, there are many well-characterized Gal4 enhancer lines that lack cognate LexA lines. Here, we adopted a CRISPR/Cas9-mediated gene editing approach (Lin and Potter 2016) to develop a genetic conversion system to replace Gal4 with LexA in unique Gal4-based enhancer trap lines.

Compared to existing LexA-expressing lines, the number of fly lines harboring a LexAop-regulated transgene is even smaller, and unlike the Gal4/UAS system, there is no public fly strain resource for LexAop-based shRNA expression. Thus, when a shRNA is proven to be highly specific and robust in generating phenotypes by the Gal4/UAS system, there should be a simplified cloning step to generate a counterpart in LexA/LexAop system. Here we developed a streamlined workflow to move functionally validated shRNA sequences from UAS-based transgenic lines to LexAop-based transgenic lines. To expand the development of these genetic tools and methods, we created secondary school and undergraduate courses in fly transgenesis. From student researchers and instructors comprising an international scholastic network, we generated novel LexA and LexAop-based fly lines permitting tissue-specific expression of shRNAs, and functionally characterized a subset of these in wing development.

## RESULTS

### Generation of LexAop-based transgenic flies for shRNA expression

To develop LexAop-dependent suppression of selected genes, we constructed a plasmid vector pWALEXA20 (White-AttB-LexAop: **Figure 1A**) permitting expression of short hairpin RNAs (shRNAs) from sulA-derived LexA DNA-binding motifs (LexAop2: Pfeiffer et al 2010). The vector design and use of universal polymerase chain reaction (PCR) primers enables insertion of validated shRNA sequences from the Transgenic RNAi Project 3 collection (TRiP-3: Ni et al 2011) adjacent to multimerized LexAop elements. This streamlined cloning process has been successfully adopted in U.S. secondary schools, and has facilitated the construction of multiple TRiP-3 shRNA-based lines (**Methods: Figure 1B**).

**Figure 1.**
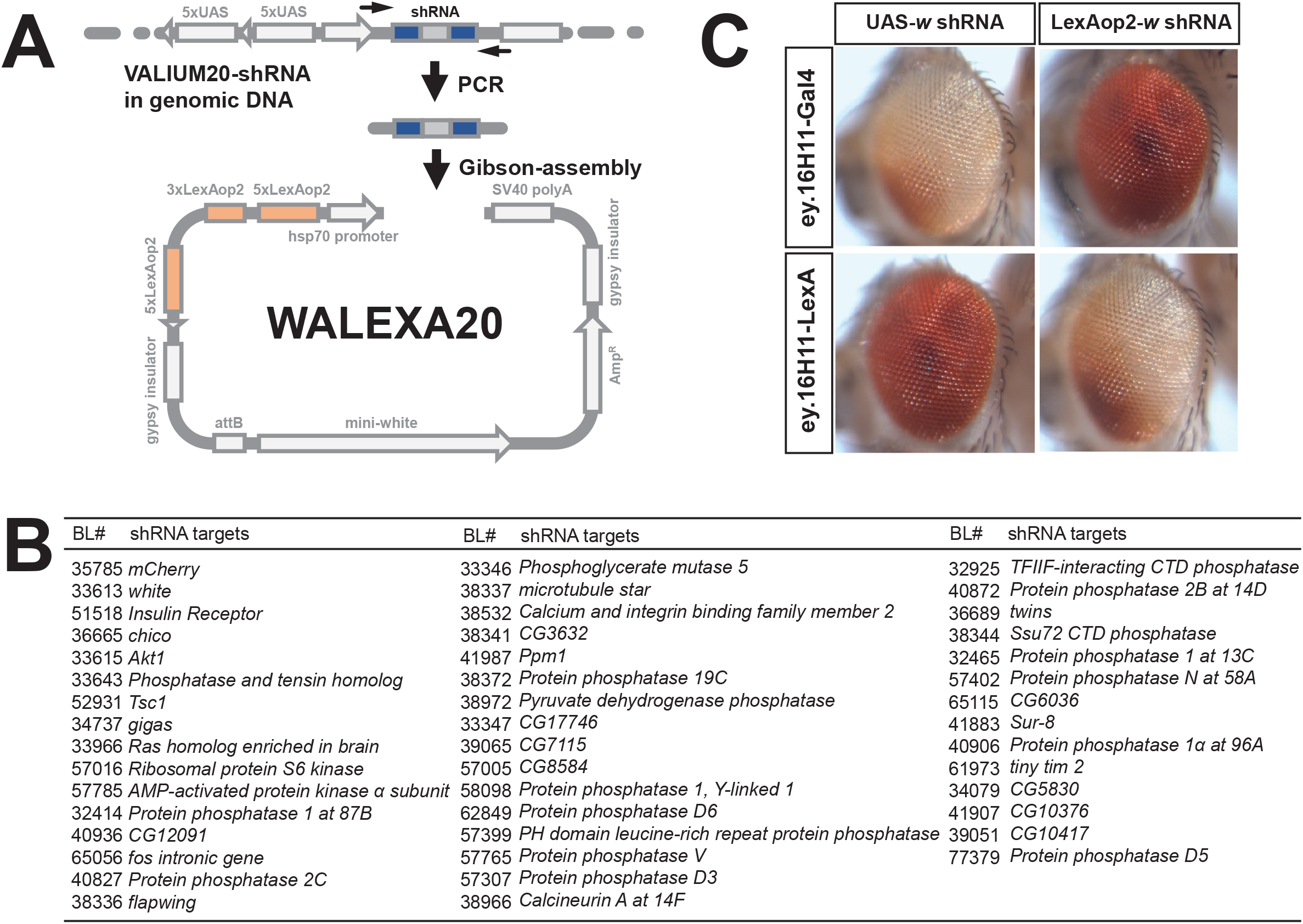
Graphical summary of cloning gene-specific shRNAs to a LexAop2-based shRNA expression vector and functional verification of LexAop-based shRNA system in adult eyes. (A) Gene-specific shRNA sequence (blue) was amplified from genomic DNA of VALIUM20-based transgenic flies using universal primer pairs (black arrows). PCR products were directly used in Gibson-Assembly reaction to clone shRNA sequences (blue) to pWALEXA20 vectors harboring LexAop2 enhancers (orange). (B) List of target genes for VALIUM20-based UAS-shRNA transgenic lines and their identifiers in Bloomington Drosophila Stock Center (BL#). shRNA sequences of the target genes are listed in Table S1. (C) Functional comparison of UAS- and LexAop-based shRNA transgenic lines targeting the *white* gene in adult eyes. Adult eyes are oriented anterior to the left. Eyeless enhancers (16H11) are more effective in knocking down *white* gene expression in the dorsal posterior (white eye area in the top-left and bottom-right panels) than ventral anterior areas of the eye in Gal4/UAS and LexA/LexAop combinations. Wild-type eye colors are maintained in Gal4/LexAop and LexA/Gal4 combinations, indicating these two systems are functionally independent.

To evaluate LexA-regulated shRNA function, we first used a transgenic fly line harboring an *eyeless* enhancer (GMR16H11) driving LexA (Pfeiffer et al 2010) to direct expression of *white* shRNA in eyes, then assessed the loss of eye pigment as a readout for *white* gene knockdown efficiency. In GMR16H11-LexA, LexAop-*white* shRNA flies, we observed a darker eye color in the ventral eye field, and lighter in the dorsal field (**Figure 1C**), indicating localized suppression of *white*. This was comparable to the pattern of eye color in GMR16H11-Gal4, UAS-*white* shRNA flies (**Figure 1C)**. Thus, tissue specific suppression of *white* was comparable in the LexA/LexAop and Gal4/UAS systems, though the degree of eye color reduction was more severe in Gal4/UAS progeny. By contrast, we observed no detectable change in eye color of control GMR16H11-Gal4, LexAop-*white* shRNA flies, or in GMR16H11-LexA, UAS-*white* shRNA flies (**Figure 1C**), providing additional evidence that LexA/LexAop and Gal4/UAS systems function independently without any cross-activation or leaky expression of shRNA.

### Gal4 to LexA conversion in an enhancer trap line by homology assisted CRISPR/Cas9 knock-in

Adult wing size and morphology have been extensively used to identify genes regulating cell growth and patterning in *Drosophila* development. Assessments of wing phenotypes for their size, shape, and vein patterns are highly reproducible and are measurements readily adoptable by early-stage scientists, including secondary school researchers (**Methods**). There are well characterized wing-specific enhancer trap Gal4 lines such as P{GawB}*bbg*^C96^ and P{GawB}*Bx*^MS1096^ (‘MS1096-Gal4’ hereafter; Capdevila and Guerrero 1994; Neumann and Cohen 1996) in which expression of the Gal4 sequence in P{GawB} is regulated by wing-specific enhancers, which remain unidentified..To generate comparable LexA lines with these unique enhancer trap Gal4 lines, we developed approaches to achieve homology assisted CRISPR knock-in (HACK: Lin and Potter 2016: **Methods**) to replace the existing Gal4-encoding DNA sequence with a LexA.G4H. To assess this approach, we targeted the MS1096-Gal4 enhancer trap line (**Figure 2A, B**). The trans-chromosomal conversion of MS1096-Gal4 target on the X chromosome to LexA.G4H using a HACK-based donor on the second chromosome was efficient (four independent conversion events in 80 individual crosses; **Figure 2**). To address the specificity of LexA expression in the converted lines, we assessed expression of a dual fluorescence reporter in the wing discs of MS1096-LexA.G4H, LexAop-tdTomato, UAS-Stinger larva. The expression of LexAop-based reporter by MS1096-LexA.G4H (**Figure 2C**, red) revealed a comparable pattern to the original MS1096-Gal4. Moreover, we did not observe cross-activation of the UAS-based reporter, indicating LexA/LexAop and Gal4/UAS systems function independently. In summary, we generated fly lines permitting CRISPR/Cas9-mediated conversion of Gal4-to LexA-expressing lines, demonstrated the feasibility and efficiency of this genetic conversion, and confirmed the maintenance of the tissue-specific LexA expression in the LexA-converted progeny.

**Figure 2.**
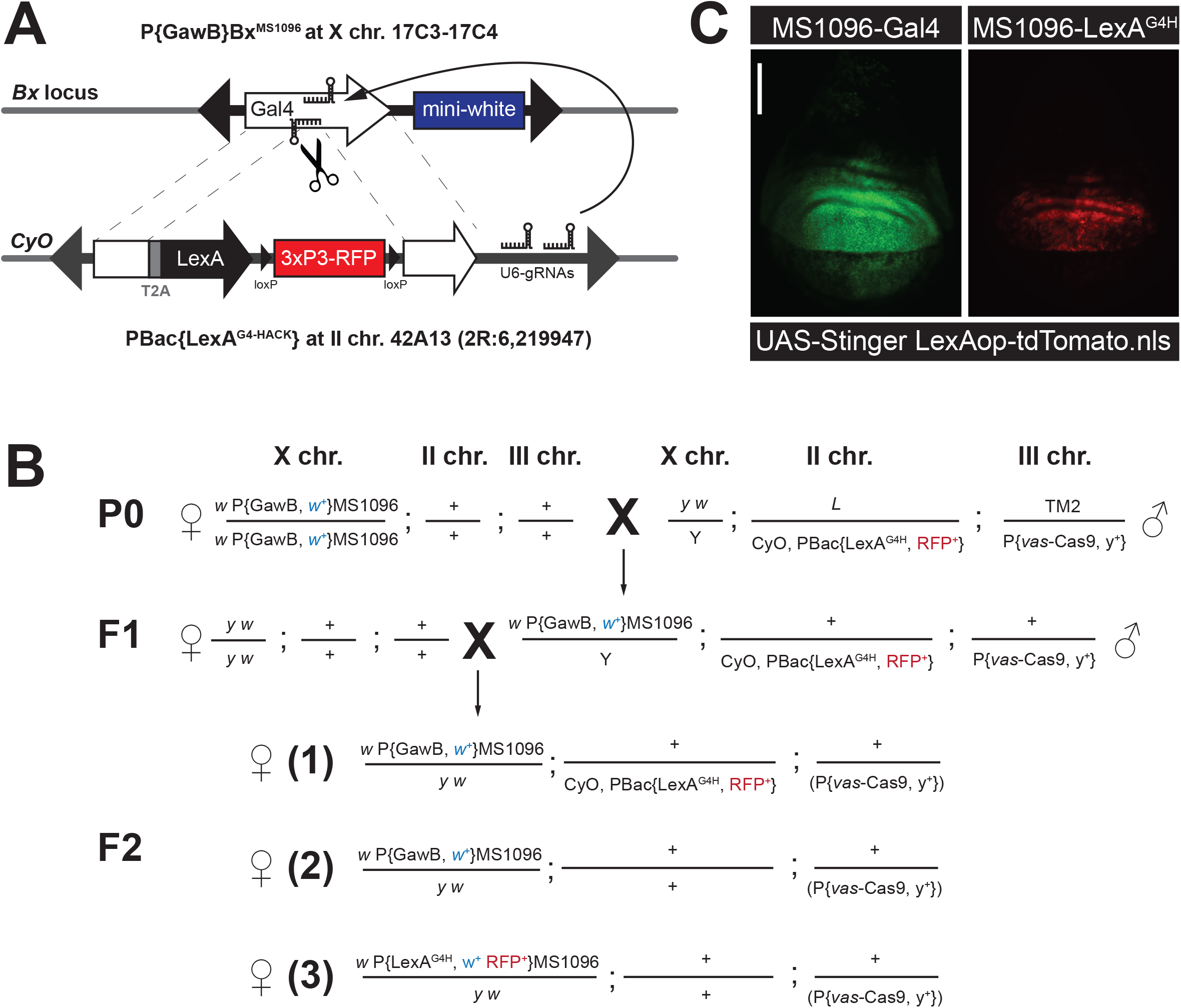
Schematic outlines of trans-chromosomal Gal4 to LexA.G4HACK conversion system and genetic crosses for identifying the conversion events. (A) The LexA.G4HACK donor is located at 42A13 on the second balancer chromosome, CyO. In the germline, Cas9 (scissors) and two guide RNAs (U6-gRNAs) make double-strand breaks in the middle of Gal4 encoding sequence (the white arrow at Bx locus on the upper X chromosome), allowing it to be repaired by the donor sequence carrying Gal4 homology sequences (the white bar and arrow on the lower CyO chromosome) that are in-frame with T2A-LexA encoding sequence. Successful repair events are identified by co-segregation of both mini-white transgene eye color (blue box) and red fluorescent protein eye markers (red box, 3xP3-RFP). (B) Homozygous females for the target P{GawB} element harboring Gal4 encoding sequence and mini-white (*w*^+^) were mated to males carrying both a donor LexA.G4H transgene marked by an eye fluorescence marker 3xP3-RFP (PBac{LexA.G4H}) and a germline-specific Cas9 (P{vas-Cas9}) marked by a body-color marker *y*^+^. The tri-transgenic F1 male progeny were mated to *y*^1^ *w*^1118^ females individually. Of the different F2 possibilities (1-3), Gal4 to LexA.G4H conversion events were identified in F2 females without CyO balancer, but with mini-white eye color and RFP eye fluorescence marker (3). (C) Comparison of the original MS1096-Gal4 and converted MS1096-LexA.G4H expression by nuclear GFP (UAS-Stinger) and nuclear tdTomato (LexAop-tdTomato.nls) reporters in larval wing discs. The reporter expressions are mostly restricted in the dorsal half of the wing disc pouch area in both Gal4 and LexA.G4H driver lines.

### Functional tests of LexAop-based shRNA transgenic lines in developing wings

Insulin signaling regulates the growth of developing tissues by modulating both cell number and size (Rulifson et al 2002; reviewed in Kim et al 2021). Measurement of adult wing size has been used to quantify disruption of insulin signaling during *Drosophila* development (Park et al 2014). To assess if the new wing-specific LexA line is suitable for shRNA-mediated suppression of genes encoding insulin signaling pathway regulators, we generated fly lines harboring LexAop-shRNA transgenes targeting *Insulin Receptor* (*InR*), *chico* (an orthologue of mammalian *Insulin Receptor Substrates 1/2*), *Phosphatase and tensin homolog* (*Pten*), *Akt*, and *Ras homolog enriched in brain* (*Rheb*) (**Figure 3A**).

**Figure 3.**
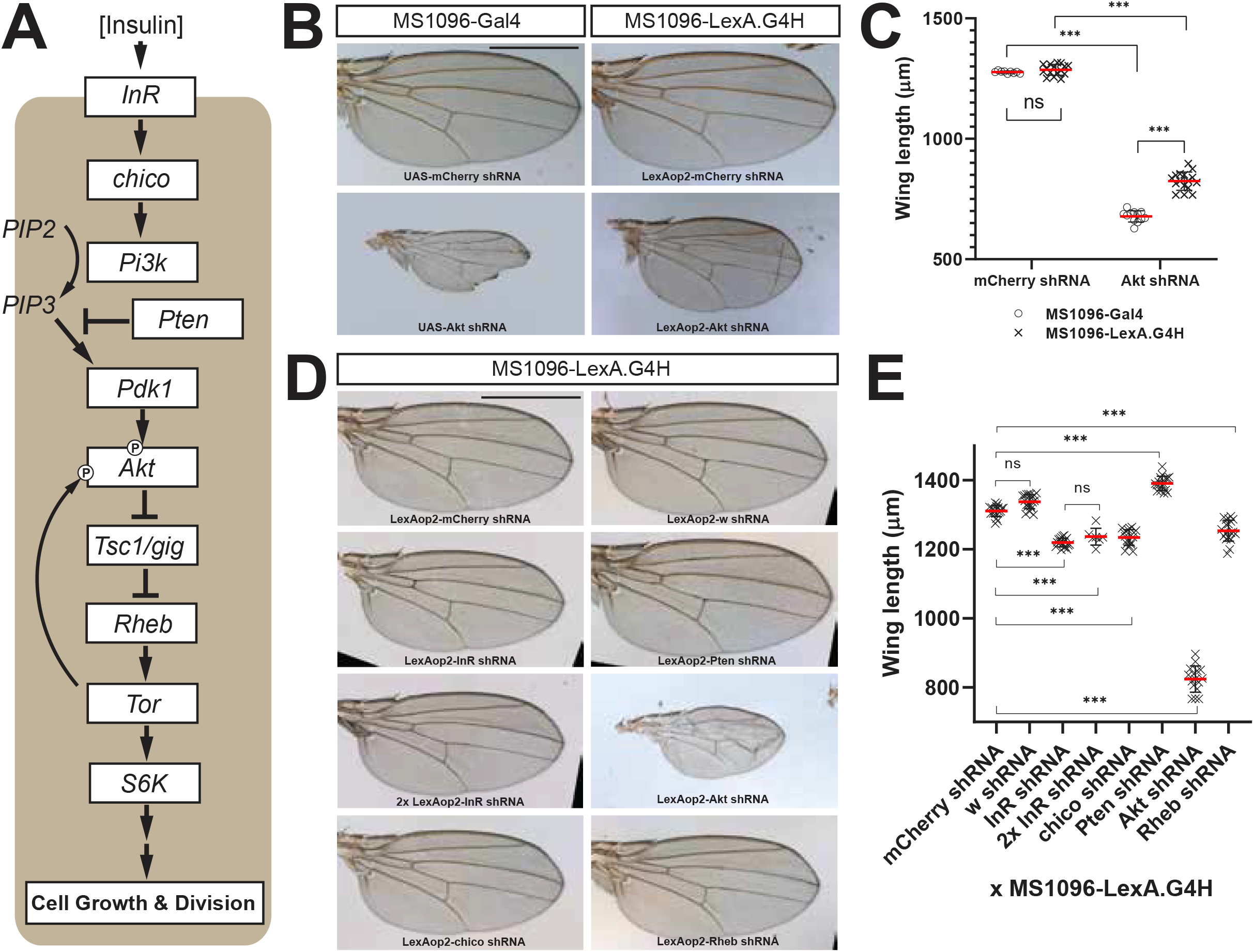
Functional validation of LexA/LexAop shRNA system in adult wings for insulin signaling regulators. (A) The insulin signaling cascade and its components regulating cell size and proliferation during animal development. The arrows indicate positive regulations between signaling components, and ‘T’ symbols between components indicate negative regulations. The two ‘P’ symbols on the Akt protein indicate different phosphorylation sites by Pdk1 and Tor. (B) Comparison of adult wings expressing control shRNA (*mCherry* shRNA) or *Akt* shRNA by Gal4/UAS or LexA/LexAop systems. Compared to *mCherry* shRNA expression, *Akt* shRNA expression resulted in smaller wing sizes in both systems. The black scale bar indicates 500 μm. (C) Quantification of wing length in animals expressing *mCherry* or *Akt* shRNAs by either Gal4/UAS or LexA/LexAop systems. *** indicates the statistical significance of P<0.001 in student’s *t*-test and ns indicates statistically not significant. The error bars are standard deviations. The red bars are the average length of n ≥ 9 wings. (D) Adult wings expressing shRNAs targeting insulin signaling component genes by LexA/LexAop system. (E) Quantification of wing length expressing shRNAs by LexA/LexAop system. Compared to *mCherry* shRNA controls, all shRNAs targeting insulin signaling component genes either increased or decreased the wing length significantly (P<0.001) while the shRNA targeting *white* gene did not (ns). Increasing the shRNA copy number did not make the wing smaller than a single copy of shRNA for the Insulin receptor gene (ns). *** indicates the statistical significance at P<0.001 in student’s *t*-test, and ns indicates statistically not significant. The error bars are standard deviations. The red bars are the average length of n ≥ 9 wings.

To compare LexAop-dependent shRNA-based gene suppression to the original UAS-driven shRNA lines, we intercrossed MS1096-LexA.G4H lines to LexAop-shRNA lines: progeny from intercrosses of the original MS1096-Gal4 and corresponding UAS-shRNA lines served as controls. In adult flies, we observed that shRNA targeting of a crucial insulin signaling gene, *Akt*, resulted in significantly smaller adult wings compared to the control shRNA targeting *mCherry* in both Gal4/UAS and LexA/LexAop systems, although wings after Gal4/UAS-based knockdown of *Akt* were significantly smaller compared to wings deploying LexA/LexAop knockdown (**Figure 3B, 3C**). In contrast, wings expressing the control *mCherry* shRNA by the two systems did not differ in size (**Figure 3B, 3C**), indicating the measurement of wing size is highly reproducible and LexA expression itself does not interfere with wing development. In addition, adult flies with wing-specific suppression of *Akt* eclosed without any discernable developmental defect, other than wing size changes. These findings are consistent with restricted expression of MS1096-LexA.G4H and MS1096-Gal4 to wing discs.

To assess the regulatory function of other genes in the insulin signaling pathway, we next assessed the impact of LexAop-shRNA lines targeting *InR, chico, Pten*, or *Rheb* on wing size. In adult flies, we observed that shRNAs targeting of *InR, chico*, and *Rheb* resulted in significantly smaller adult wings (**Figure 3D, 3E**), consistent with the known requirement of these genes for cell growth while shRNA targeting *white* whose function is not involved in insulin signaling did not change the size. In contrast, shRNA targeting of *Pten* increased wing size (**Figure 3D, 3E**), confirming the prior observation that loss of *Pten* function leads to increased eye and wing size (Goberdhan et al 1999). These results confirm the efficiency and reproducibility of the LexA/LexAop-based gene silencing strategy. To test if the degree of the observed loss-of-function phenotype can be further exerted by increasing the copy number of the LexAop-shRNA transgene, we scored the wing size of adult fly carrying one and two copies of LexAop-shRNA targeting *InR*. This effectively tested whether the LexA-driven expression levels of the shRNA, or, alternatively, the downstream cellular components of the RNA interference pathway are the limiting factor for the phenotypic strength in the wing development context. A single copy of the transgene encoding shRNA that targeted *InR* resulted in smaller wings; doubling the copy number of the same transgene did not reduce wing size further **(Figure 3D, 3E)**. These results suggest that LexA-dependent shRNA expression from one copy of the LexAop-*InR* shRNA transgene may be sufficient to saturate the capacity of the RNA interference pathway. In summary, we generated a novel fly line expressing LexA in the larval wing disc to assess the impact of LexAop-based shRNA gene suppression on wing development and growth. Generation and functional validation of LexAop-shRNA lines targeting insulin signaling motivated the development of curricula at partnering secondary schools to generate additional resources, and characterize growth regulators of wing disc development, as detailed below.

### An interscholastic network for systematic generation of new LexAop flies

To expand the repertoire of extant fly strains permitting LexA-dependent genetics, we next leveraged our multi-institutional network of secondary schools (**Methods**) to develop curricula that permit students and their instructors to generate additional new lines expressing shRNA. To assess the quality of the data generated in distant classroom settings, we assigned construction of shRNA lines targeting known insulin signaling regulators (*Tsc1, gig*, and *S6K*) in addition to the *Drosophila* phosphatases whose function have not been systematically assessed for cell growth and patterning during development. Students and instructors intercrossed these new LexAop-shRNA lines to adults expressing the wing-specific MS1096-LexA.G4H driver, then assessed adult wing length. Progeny from eight intercrosses had increased average wing length, including flies targeting *Tsc1, gig, CG17746, CG8584, Pten, Pp2B-14D, Ssu72*, and *Pp1-13C* (**purple bars, Supplementary Figure S1**) and six intercrosses targeting *S6K, Rheb, Pp1-87B, mts, PpV* and *PpD5* had reduced wing length (**green bars, Supplementary Figure S1**). The results confirm that the students correctly produced shRNA lines targeting *Tsc1, gig and S6K*, and also successfully altered wing size by targeting these genes or other known insulin signaling regulators, *Pten* and *Rheb*. Additionally, students also identified several phosphatases that altered cell growth or patterning in adult wings (**Supplementary Figure S1)**.

We repeated a subset of these intercrosses and confirmed students’ results of statistically significant increases or reductions of wing length in adult progeny from these crosses compared to *mCherry* or *white* shRNA controls (**Figure 4A**). For example, shRNAs targeting *CG17746, CG8584, Pp2B-14D, Ssu72, Pp1-13C, PpV*, and *PpD5* resulted in wing size changes without any patterning defect assessed by wing hairs and vein locations: further studies should determine if these phosphatases are directly or indirectly involved in the insulin signaling pathway. By contrast, the expression of *Pp1-87B* shRNA or *mts* shRNA led to the development of morphologically abnormal wings (**Figure 4B**). *Pp1-87B* and *mts* genes are previously shown to regulate mitotic chromosomal segregation, Hedgehog signaling, and Wingless signaling (Chen et al 2007; Luo et al 2007; Zhang et al 2009). To understand the basis of dysmorphic wing development after shRNA-mediated *Pp1-87B* or *mts* gene knockdown, we immunostained wing discs to assess an earlier stage of wing development in third instar larvae. Expression of phospho-Histone H3, a cell cycle marker of G2/M2 transition, was not detectably altered after *Pp1-87B* or *mts* gene knockdown (**Figure 4B**). However, we observed increased expression of Cleaved Caspase 3, a marker of apoptosis, in the wing disc cells expressing *Pp1-87B* shRNA or *mts* shRNA (**Figure 4B**). These results suggest that the dysmorphic adult wing phenotypes after *Pp1-87B* or *mts* gene knockdown may originate from a significant increase of cell apoptosis during wing development. The increased expression of Cleaved Caspase 3 in wing discs expressing *Pp1-87B* or *mts* shRNAs was restricted to the central region of the wing disc where the expression of our MS1096-LexA.G4H is highest, providing additional evidence for the tissue specificity of the new LexA/LexAop-based gene knockdown system. Thus, classroom-based experimental studies revealed several phosphatases that may be required for sustaining cell growth and survival during wing development.

**Figure 4.**
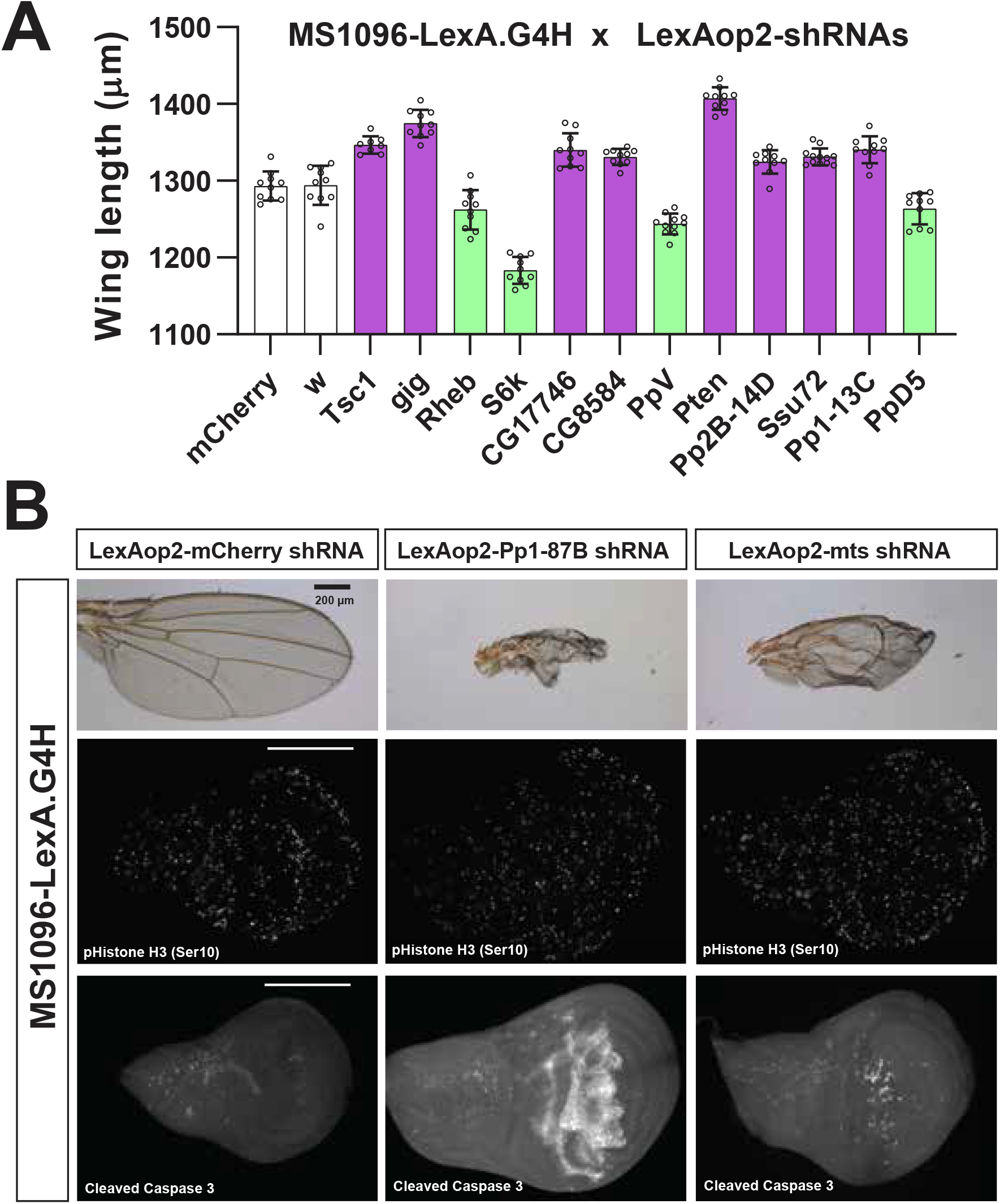
Identification of growth regulators for Drosophila wing development using the LexA/LexAop system. (A) Quantification of wing length in flies expressing selected shRNAs that resulted in significantly smaller (green bars) or larger (purple bars) wings compared to the control *mCherry* shRNA expressing wings (P<0.01, student’s *t*-test). The error bars are standard deviations. The average lengths are based on measurements of 8 individual female wings in each genotype. (B) Altered adult wing morphology and increased number of apoptotic cells in larval wing discs expressing *Pp1-87B* or *mts* shRNAs compared to the control *mCherry* shRNA. Cleaved caspase 3 staining marks apoptotic cells while phospho-Histone H3 staining marks proliferating cells. *PP1-87B* and *mts* shRNA expression by MS1096-LexA.G4H resulted in increased apoptotic markers in the dorsal wing pouch where MS1096 enhancer expression is high **(Figure 2C)**, but the marker expressions are not changed outside of LexA.G4H expression domain in wing discs.

## DISCUSSION

Intersectional binary expression approaches such as simultaneous use of the LexA/LexAop and Gal4/UAS systems have empowered fly biology, particularly in studies of neuroscience, and intercellular communication coordinating metazoan metabolism (Kim et al 2021). For example, simultaneous use of two independent binary expression systems allowed clonal analysis of multiple cell populations (Lai and Lee 2006; Bosch et al 2015), studies of tissue epistasis (Yagi et al 2010), and elucidation of otherwise undetected contacts between cells (Gordon and Scott 2009; Macpherson et al 2015). To address a growing need for unique fly strains permitting flexible intersectional approaches, we are expanding our efforts to create new genetic tools suitable for this purpose (Kockel et al 2016, 2019; Wendler et al 2020). Here we describe (1) a genetic tool to convert existing Gal4 lines to LexA.G4H by CRISPR/Cas9-mediated gene editing, (2) a universal cloning method to generate LexAop-based shRNA transgenic lines from functionally validated UAS-based shRNA transgenic lines, and (3) a research-based pedagogy to leverage effort by collaborating students in our secondary school network (Kockel et al 2016, 2019). Our results demonstrate that *in vivo* gene regulation by LexAop-based gene suppression succeeded in multiple tissue contexts. We also showed the functional independence of the LexA/LexAop and Gal4/UAS systems, and thus the potential for their simultaneous use within independent cell types. The resources reported here can be applied to generate additional intersectional binary expression tools in studying genetics, development, physiology, metabolism, and other studies of cells and tissues.

An analysis of HACK-based conversion of Gal4 to QF2.G4H (Lin and Potter, 2016) showed that some target locations had higher conversion rates (conversion ‘hot spots’) while most donor locations worked equally well for intra-chromosomal or trans-chromosomal conversions. This led us to place our LexA.G4H donor transgene on the second chromosome balancer, CyO, so that the parental donor transgene can be easily identifiable in the F2 progeny while the conversion events at the target location can be detected efficiently. Using this CyO-bound donor transgene on the second chromosome, the trans-chromosomal conversion of MS1096-Gal4 target on the X chromosome to LexA.G4H was successful and efficient. Although the conversion rates could vary for other target locations, our donor transgene located on the CyO chromosome may be sufficient for the general purpose of establishing LexA.G4H lines on other chromosomes. In this regard, additional Gal4 lines on the second and third chromosomes have been successfully converted to LexA.G4H through the interscholastic network, and characterization of these new LexA.G4H lines is in progress (T. Chisholm, A.E. Rankin, S. Kim and S. Park, unpublished results).

To address the challenge of validating the specificity and efficiency of gene suppression using shRNA approaches *in vivo*, we sought to generate a facile cloning step to develop genetic counterparts in the LexA/LexAop system for specific shRNAs that have proven to be robust in the Gal4/UAS system. Here, we designed universal primers that bind immediately outside of the UAS-shRNA region in transgenic flies from the TRiP-3 collection, then streamlined the production of strains harboring functionally validated shRNA sequences expressed from LexAop2 sequences. While characterizing LexAop-based shRNA lines in adult eyes and wings, we consistently observed slightly weaker shRNA activities in LexA/LexAop system compared to the corresponding Gal4/UAS system. Given that both systems share identical transactivation domains, this difference could reflect lower affinity of the LexA DNA binding domain to multimerized (13x) LexAop2 sequences in the pWALEXA20 vector. If so, the shRNA expression system may be further refined by substituting for 8x, 16x, or 26x LexAop2 sequences to decrease or increase the expression level of transgenes (Pfeiffer et al 2010). Alternatively, the activity of the LexA-encoding sequence used in this study (nlsLexA::GADfl, **Methods**) may be further optimized (Yagi et al 2010). In summary, new approaches presented here generated the wing-specific LexA driver line, MS1096-LexA.G4H, and LexAop-based shRNA lines that augment the arsenal of available LexA-dependent expression tools (Pfeiffer et al 2010; Kockel et al 2016) and also complement existing Gal4/UAS resources.

Data and biological resources here were generated from partnerships connecting research universities with teachers and students at secondary schools. This illustrates the feasibility of building an interscholastic network to generate unfulfilled scientific resource needs, while providing paradigms for STEM education. Resources and outcomes described here significantly extend, develop and complement the interscholastic partnerships in experiment-based science pedagogy described in our prior studies, which focused on generating LexA enhancer trap lines (Kockel et al 2016, 2019). The current study involved collaborations between Stanford and Oxford University investigators with students and teachers at the Lawrenceville School and the Phillips Exeter Academy, two U.S. secondary schools. Together, we developed courses focused on shRNA cloning and transgenesis, or CRISPR/Cas9-mediated conversion of Gal4 to LexA strains. These research experiences for students include generating novel fly strains, accompanied by a sense of scientific discovery and ownership (Hatfull et al 2006), and connection to a broader community of science through the distribution of flies, information, and resources through stock centers and scientific meetings. A curriculum based on classical and molecular fruit fly genetics combined with developmental biology provided a compelling framework of authentic research experiences for student scientists in U.S. grades 9-12 (Kockel et al 2016, 2019; Redfield, 2012). The success of this experience affirms the value of experience-based practice in secondary education, and we speculate such benefits may be amplified from even earlier introduction of experimental science in the U.S. K-12 sequence. In summary, this experience demonstrates how longitudinal studies involving molecular biology, genetics, and developmental biology can build a thriving, interconnected network of teachers, students, and classes that impacts personal growth, and professional development, and in the process generates valuable resources and data for the global scientific community.

## METHODS

### *Drosophila* strains

Except for transgenic lines that were generated in this study, all other Drosophila lines were obtained from the Bloomington Drosophila Stock Center.

### Construction of pWALEXA20 (<u>W</u>hite-<u>A</u>ttB-<u>LexA</u>op) vector

To replace 10xUAS in the pWALIUM20 vector with 13xLexAop2 (*sulA* derived LexA DNA-binding motifs), a 675 bp product was amplified from pJFRC19-13xLexAop2-IVS-CD4-myr::GFP (Pfeiffer et al 2010) using primers LexAop_F1 (5’-CACCCATGCATAGGGCCGCAAGCTTGCATG-3’) and hsp70p_R (5’-CCTTTTAGATCTATTCAGAGTTCTCTTCTTGTATTCAATAATTACTTCTTGGC-3’). The PCR product was inserted to the PstI and BglII sites on the pWALIUM20 vector (Perkins 2015) by Gibson-assembly (NEB HiFi DNA Assembly Kit) to generate pWALEXA20 (White-AttB-LexAop for shRNA expression), a phiC31 integrase-mediated transformation vector with mini-white as a selection marker.

### Cloning of shRNA sequences from genomic DNAs of VALIUM20 based TRiP-3 lines

0.5 μg of pWALEXA20 vector was digested with XbaI and NdeI and run on 0.8% agarose gel in 1x Tris-Acetate EDTA (TAE). The resulting 9947 bp fragment was purified in 10 μl water using the Zymoclean Gel DNA Recovery Kit.

To extract genomic DNA from shRNA-based Transgenic RNAi Project (TRiP-3) transgenic lines, 1 male fly was ground in 50 μl of Squishing Buffer (10 mM Tris-HCl pH8, 1 mM EDTA, 25 mM NaCl, and 0.2 mg/ml Proteinase K). The sample was incubated at 37°C for 30 minutes, heat-inactivated at 95°C for 3 minutes, and then centrifuged at >18,000 g for 5 minutes. 1 μl of the clear supernatant containing the extracted genomic DNA was added to 19 μl of PCR master mix containing 7 μl of water, 10 μl of Q5 Hot Start High-Fidelity 2x Master Mix (NEB M0494S), 1 μl of 10 μM shRNA_GA_F primer (5’ GAGAACTCTGAATAGATCTGTTCTAGAAAACATCCCATAAAACATCCCATATTCA-3’), and 1 μl of 10 μM shRNA_GA_R1 primer (5’-CTCTAGTCCTAGGTGCATATGTCCACTCTAGTA-3’). 1 μl of Squishing Buffer was added to 19 μl of PCR master mix as a control reaction. After a 30-second denaturing period at 98°C, 40 cycles of PCR amplification were performed with a 10-second denaturing period at 98°C, a 20-second annealing period at 62°C, and a 30-second extension period at 72°C. 5 μl of the PCR reaction was then run on 3% agarose gel in 1xTAE to confirm the 202 bp PCR product. 49 VALIUM20 based TRiP-3 lines from the Bloomington Drosophila Stock Center were processed as above, and we successfully amplified the shRNA region in all but three lines (Bloomington Stock ID 43283, 57215, and 57519), resulting in a 93.8% cloning success rate. Obtaining replacement lines for the three lines that went uncaptured did not solve the problem.

Once PCR amplification was confirmed, 1 μl of the PCR reaction was added to 9 μl of Gibson-assembly reaction containing 3.5 μl of water, 5 μl of NEBuilder HiFi DNA Assembly Master Mix, and 0.5 μl of the linearized and purified pWALEXA20 vector. 1 μl of water was added to 9 μl of Gibson-assembly reaction as a control reaction. After incubating the Gibson-assembly reaction at 50°C for 1 hour, 25 μl of competent cells (NEB E5520S) were transformed with 1 μl of the reaction, and all cells were plated on LB agar plate with ampicillin using L-shaped bacterial cell spreader.

Bacterial colonies were picked and grown in 10 ml LB broth with ampicillin for 16 hours at 37°C. Amplified plasmids were purified by Zyppy Plasmid Miniprep (Zymo Research) and sequenced from the *hsp70* basal promoter using the primer hsp70p_F (‘5-CTGCCAAGAAGTAATTATTGAATACAAGA-3’) to confirm successful cloning of shRNA fragments.

All cloned shRNA sequences in the pWALEXA20 vector were identical to the reported sequences except two sequences (originated from Bloomington Stock # 33613 for *white* and 41883 for *Sur-8*) in which there were single base pair mismatches within the complementary region **(Table S1)**. We found that identical mismatches were also present in the genomic DNA of the originated stocks. We do not know when these changes were introduced to these stocks, but we do not expect these changes would affect their ability to knock down their target genes since the *white* shRNA sequence with a single mismatch was shown to be functionally active **(Figure 1C)**.

### Generation of transgenic LexAop2-shRNA lines

All sequence-verified plasmids were injected to either y^1^ w* P{nos-phiC31}; P{CaryP}attP40 (#79604) or y^1^ w* P{nos-phiC31}; P{CaryP}attP2 for integration into the second or third chromosome, respectively. To verify the sequences of the newly generated transgenic lines, 232 bp PCR products were amplified from genomic DNA using hsp70p_F and shRNA_GA_R1 primers, purified using DNA Clean and Concentrator-5 (Zymo Research), then sequenced using shRNA_GA_R1 primer.

### Functional test of *white* gene knockdowns in adult eyes

To knockdown *white* gene expression in adult eyes, P{y[+t7.7] w[+mC]=GMR16H11-Gal4}attP2 (BL#47473) or P{y[+t7.7] w[+mC]=GMR16H11-lexA}attP40 (BL#61516) was intercrossed to either y[1] v[1]; P{y[+t7.7] v[+t1.8]=TRiP.HMS00004}attP2/TM3, Sb[1] (BL#33613) or y[1] v[1]; P{y[+t7.7] w[+mC]=LexAop2.HMS00004}attP40 (this study). 5-day old F1 female eyes were imaged by QCapture software (Quantitative Imaging Cooperation).

### Generation of MS1096-LexA.G4H line

The second chromosome balancer, CyO, carrying a PBac{y^+^ = attP-9A} was generated by mobilizing PBac{y^+^ = attP-9A} VK00006 on the X chromosome to the chromosomal location 2R (42A13) and the molecular location at 2R:6,219,947 on the CyO. Using the primers LexA_HACK_F1 (5’-AGAACCCCGGGCCCCCTAGGATGCCACCCAAGAAGAAGCG-3’) and LexA_HACK_R1 (5’-TGAATAATTTTCTATTTGGCTTTAGTCGACGGTATCGATAAG-3’), 2926 bp fragment was amplified from the template pBPnlsLexA::GADflUw vector (Addgene #26232) and the amplified product was assembled to 9083 bp fragment of pHACK-Gal4>QF2 (Addgene #104873) that was digested with AvrII and SalI. The resulting construct called pHACK-Gal4>nlsLexA::GADfl was inserted to the PBac{y^+^ = attP-9A} site located in the CyO chromosome. The resulting donor transgenic line, called ‘LexA.G4H’ for brevity, was combined to the vas-Cas9 transgene in the presence of L^1^ and TM2 to distinguish the chromosomes by visible markers in a y^1^ w^1118^ genetic background. w^1118^ P{w^+^ = GawB}Bx^MS1096^ (Bloomington stock #8860) females were crossed to y^1^ w^1118^; L^1^/CyO, PBac{y^+^ RFP^+^ = LexA.G4H}; TM2/P{y^+^ = vas-Cas9}VK00027 males. F1 male progeny were crossed to y^1^ w^1118^ females individually, and about 50 Non-CyO F2 virgin females were screened for w^+^ and RFP^+^ eye markers per cross **(Figure 2B)**. 4 independent MS1096-LexA.G4H lines were established from 80 individual F1 crosses and the lines were functionally tested by overexpressing LexAop2-*Akt* shRNA in wings. Female wings were mounted as described previously (Park et al 2014) and imaged in AxioImager microscope (Zeiss). Wing lengths were measured using an AxioVison microscope (Zeiss) and ImageJ software (NIH).

### Reporter expression and Immunofluorescence on larval wing discs

To visualize the expression patterns of MS1096-Gal4 and MS1096-LexA.G4H in the third instar wing discs, males of a dual reporter line, UAS-Stinger LexAop-tdTomato.nls (Bloomington stock #66680) were crossed to females of either MS1096-Gal4 or MS1096-LexA.G4H. Wing discs from F1 progeny were fixed in 4% paraformaldehyde, 1x PBS, and 0.1% Triton X-100 for 1 hour, then mounted in Antifade mounting medium with DAPI (Vectashield H-1200). Both GFP and RFP channel images were acquired and overlaid in AxioVision to visually check for any leaky interactions between Gal4/UAS and LexA/LexAop expression systems.

To assess cell proliferation and apoptosis in larval wing discs expressing *Pp1-87B* or *mts* shRNA, females of MS1096-LexA.G4H were crossed to males harboring LexAop2-shRNA transgenes. Wing discs from F1 progeny were fixed as described above, then permeabilized in PBS with 1% Triton X-100 for 1 hour. Wing discs were incubated at 4°C for 16 hours in PBS with 0.1% Triton X-100 containing either anti-phospho-Histone H3 (Ser10) antibody (Cell Signaling Technology #9701, 1:1000) for a proliferation marker or anti-cleaved caspase-3 (Asp175) antibody (Cell Signaling Technology #9661, 1:1000) for an apoptosis marker. After washing three times in PBS with 0.1% Triton X-100 for 10 minutes, the discs were incubated at 22°C for 1 hour in PBS with 0.1% Triton X-100 containing anti-rabbit Alexa Fluor 488 (Invitrogen #A11034). After washing three times again, discs were mounted and imaged as described above.

### Secondary school class and university partnerships

To facilitate the generation of new LexA and LexAop strains, we formed scholastic collaborations between Stanford University investigators and two secondary schools in the U.S. (Lawrenceville School, and Phillips Exeter Academy), and with the University of Oxford Sir William Dunn School of Pathology. Briefly, course work at Lawrenceville for the ‘Hutchins Scholars Program’ summer term was developed by Stanford, Oxford, Lawrenceville, and Exeter instructors, with the goal of generating WALEXA20-based constructs harboring validated shRNA’s amplified from relevant TRiP-3 stocks (see above). This course included instruction in DNA amplification and cloning, fly transgenesis, multigeneration intercrosses to generate genetically ‘balanced’ stocks, larval and adult dissection, tissue mounting, wing morphometry, and tissue immunohistology. Alongside the protocols taught, students were introduced to fundamental concepts in cell and developmental biology utilized, such as the central dogma and gene expression systems. To develop a separate course (Bio670) at Exeter with the goal of generating new LexA-expressing fly strains, instructors and staff at Stanford worked with Exeter instructors to develop HACK-based CRISPR/Cas9 replacement of Gal4 with LexA sequences. The instruction manuals, additional manuals for teachers, schedules and related assessment materials for students are available upon request or posted online (https://stan-x.org).

## ACKNOWLEDGMENTS

We thank the Bloomington Drosophila Stock Center for transgenic fly stocks used in this study. We thank past and current members of the Kim lab for helpful discussions and welcoming summer term students. We thank S. Murray, I. Saxe, G. Hannon, M. Rupert and R. McGuire (Lawrenceville School), A. Hobbie, E.G. Blanco, R. Weatherspoon, and W. Rawson (Phillips Exeter Academy, Access Exeter), and S. Sodywola and Montag House in the Bing Overseas Program, Oxford, for their advice, support, and encouragement. We are grateful to Philip Weissman (Micro-Optics Precision Instruments, NY) and Ken Fry (Genesee Scientific, CA) for their generous support of equipment procurement for this project. We thank Glenn and Debbie Hutchins, and the Hutchins Family Foundation, for supporting opportunities for students to engage in innovative science research at the Lawrenceville School. E.W, J.F.F, and A.S.Y.T were members of the Hutchins Scholars Program at the Lawrenceville School. Work at Phillips Exeter Academy was supported by the John and Eileen Hessel Fund for Innovation in Science Education. K.R.C. was supported by Stanford Vice-Provost for Undergraduate Education (VPUE) and Stanford Bio-X scholarships. E.G. was supported by National Science Foundation (DGE-1656518) and Institutional Training Grant in Genome Science (5T32HG000044). Work in the Kim group was supported by NIH awards (R01 DK107507; R01 DK108817; U01 DK123743; P30 DK116074 to S.K.K.), the H.L. Snyder Foundation, the Elser Trust, a Stanford VPUE faculty award, gifts from Mr. Richard Hook and from two anonymous donors, and the Stanford Diabetes Research Center.

## SUPPLEMENTARY INFORMATION

**Supplementary Figure S1.**
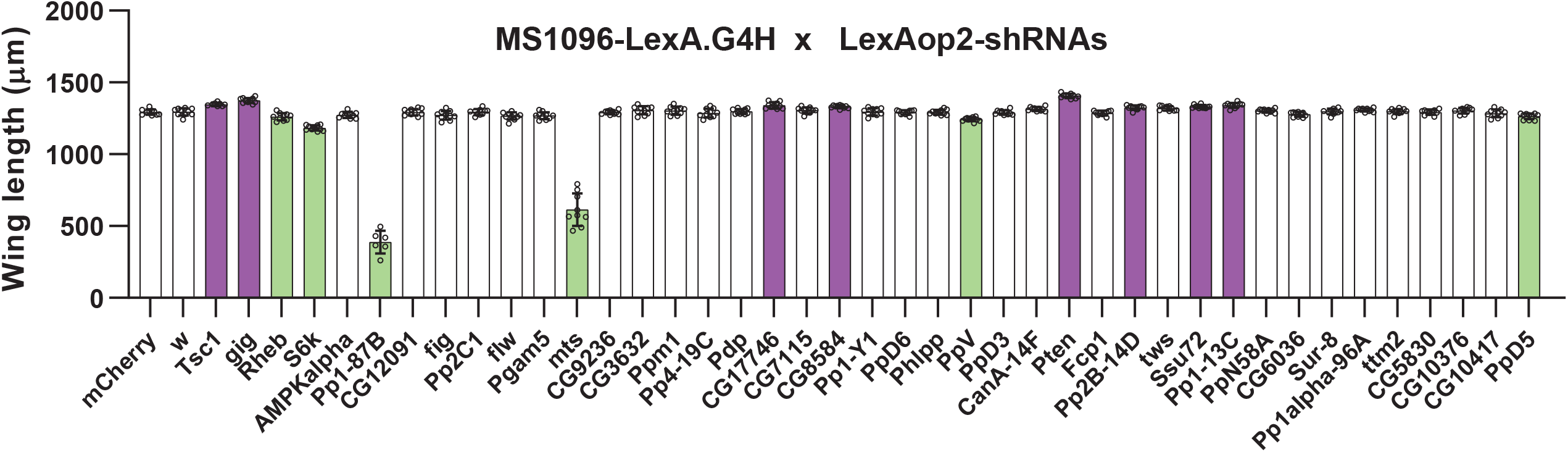
A genetic screen of growth regulators for Drosophila wing development by LexA/LexAop system. Quantification of wing length in flies expressing shRNAs for protein phosphatases and additional insulin signaling component genes. The purple bars indicate that gene-specific shRNA expression increased wing length compared to the control *mCherry* shRNA while the green bars designate decreased wing length (P<0.01, student’s *t*-test). The error bars are standard deviations. The average lengths are based on n = 8 wings in each genotype of female progeny from the indicated crosses above.

**Supplementary Table S1.**
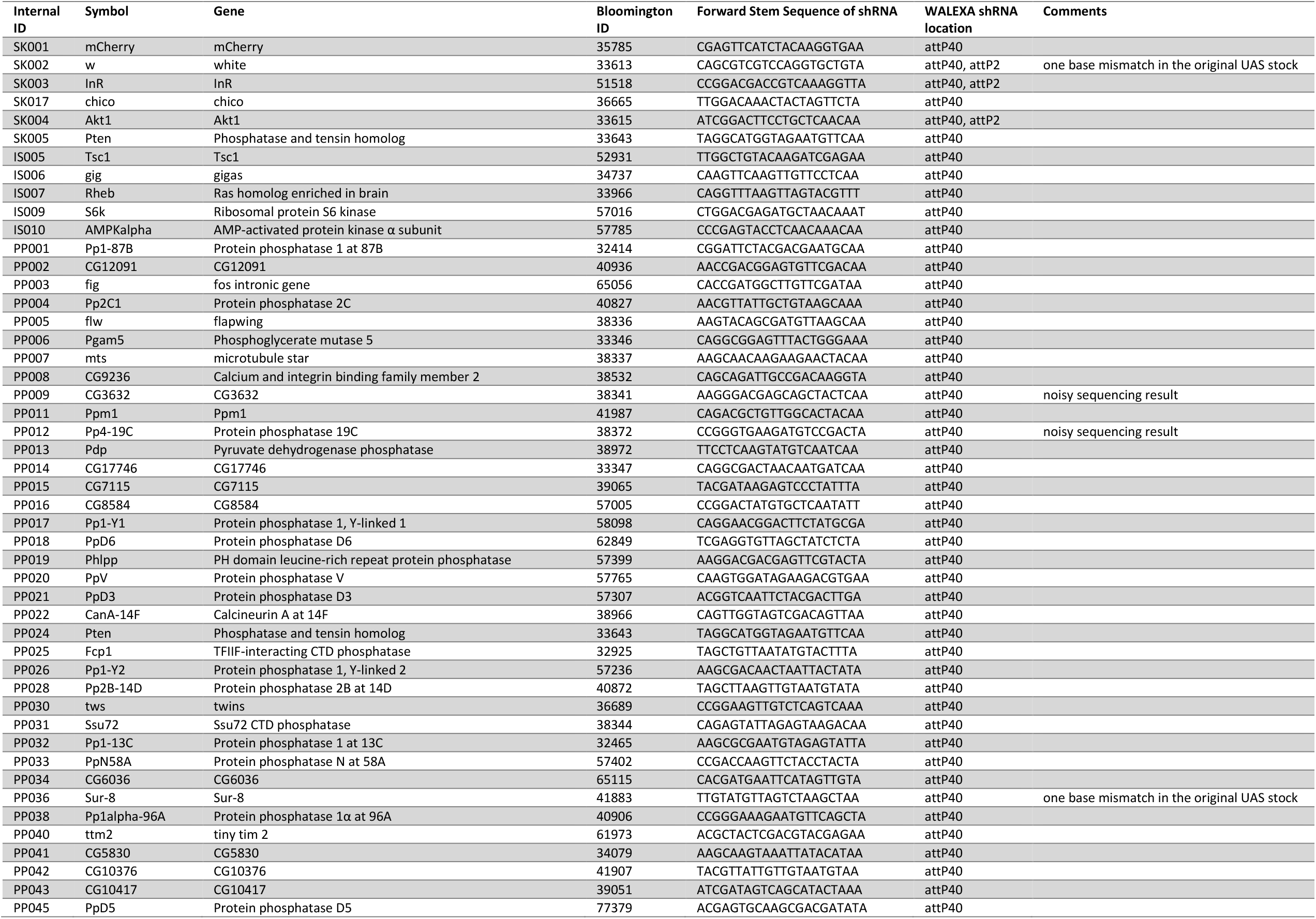
The list of genes and their LexAop-based shRNA transgenic lines generated in this study. Gene-specific shRNAs were amplified from UAS-based shRNA transgenic lines (Bloomington Drosophila Stock Center #) and the forward stem sequences of shRNAs were used to confirm the LexAop-based shRNA plasmids and transgenic lines.

